# Declining autophagy drives age-related axonal decay through microtubule alterations

**DOI:** 10.64898/2026.05.29.728680

**Authors:** Kriti Gupta, Haifa Alhadyian, Ceryce Collie, Emilia Gregory, Amaya Malmalabaduge, Natalia Sanchez-Soriano

**Affiliations:** Department of Biochemistry, Cell and Systems Biology, Institute of Systems, Molecular & Integrative Biology, University of Liverpool, Liverpool, United Kingdom; School of Molecular Biosciences, University of Glasgow, Glasgow, United Kingdom

**Keywords:** *Drosophila*, microtubules, ageing, axons, autophagy, brain, neurons

## Abstract

The microtubule cytoskeleton plays a pivotal role in maintaining axonal integrity, and its deterioration has been closely associated with neuronal decay during ageing. Autophagy, a crucial cellular degradation process, is essential for sustaining neuronal homeostasis and its dysregulation has been implicated in various neurodegenerative diseases. Changes in autophagy are also associated with ageing. However, the mechanisms by which autophagy dysfunction contributes to neuronal atrophy in the ageing brain remain poorly understood.

In this study, we unveil the significant involvement of autophagy in preserving axonal architectural integrity during ageing through regulation of the microtubule cytoskeleton. Our findings indicate that autophagic activity declines in neurons of the aged brain, and the inhibition of autophagy exacerbates age-related alterations in axonal and synaptic microtubules. This dysfunction correlates with an increase in established hallmarks of ageing, including swellings and thinning of axons plus synaptic terminal breakdown. Conversely, pharmacological and genetic strategies to enhance autophagic activity not only rescued age-related microtubule changes, but also inhibited the formation of axonal swellings, the thinning of axons and the deterioration of synaptic terminals.

Mechanistically, we find that reactive oxygen species (ROS), a consequence of impaired autophagy, lead to alterations in microtubule networks. In addition, enhancing Eb1 expression can counteract the detrimental effects of deficient autophagy, effectively blocking axonal and synaptic microtubule deterioration. These findings underscore that alterations in autophagy are a critical driver of axonal deterioration during ageing, operating through modifications to the microtubule cytoskeleton as a downstream target. Our findings deliver a cascade of events that underlie neuronal ageing.

## Introduction

Ageing profoundly impacts the delicate equilibrium of neuronal cell biology, rendering these cells increasingly vulnerable to the onset of neurodegenerative diseases (Niccoli & Partridge, 2012; Salvadores et al., 2017). Because most neurodegeneration cases are sporadic, lacking causal links to specific genetic mutations, their characteristic late onset establishes advanced age as the primary risk factor. Understanding how the ageing process undermines cellular resiliency is therefore of vital importance.

Within the adult brain, neurons face numerous challenges in maintaining the quality of their axons, dendrites, and synapses, as most neurons originate during embryonic development and must persist throughout the organism’s lifespan due to their post-mitotic nature, which can last a century in humans (Alonso et al., 2024; Sorrells et al., 2018). Due to their delicate nature, axons are particularly vulnerable; they can extend over a meter in humans and are subject to high metabolic demand (Madrer et al., 2025; Smith et al., 2023). These factors, combined with the longevity of neurons, lead to wear and tear of cellular components including organelles, the cytoskeleton and proteins in general, rendering them increasingly vulnerable to both environmental and genetic stressors and increasing their risk of developing various pathologies. These internal changes are accompanied by gradual degradative remodelling during ageing involving changes to their axons, dendrites, and synapses. An increasing body of research suggests the microtubule cytoskeleton to be central to this age-dependent remodelling (Cash et al., 2003; Fiala et al., 2007; Filipek et al., 2008; Nag et al., 2020; Niewiadomska & Baksalerska-Pazera, 2003; Okenve-Ramos et al., 2024; Shields et al., 2025; Zhang et al., 2012).

In axons, parallel bundles of microtubules are required for sustained neuronal function and maintenance by providing structural resistance and acting as tracks to transport organelles and other cellular components (Guedes-Dias & Holzbaur, 2019; Prokop, 2020). Microtubule decay is characteristic of the ageing brain. Accordingly, microtubule density and organisation are reported to be altered in ageing axons of brains across species including humans, other primates and *Drosophila* (Cash et al., 2003; Fiala et al., 2007; Okenve-Ramos et al., 2024; Shields et al., 2025). Microtubule dysfunction is associated not only with the ageing brain but also with numerous age-related neurodegenerative disorders such as Alzheimer’s disease (AD), Parkinson’s disease (PD), Amyotrophic Lateral Sclerosis (ALS), and Frontotemporal Dementia (FTD) (Cash et al., 2003; Cyske et al., 2023; Ezzo & Etienne-Manneville, 2025; Prokop, 2021; Zhang et al., 2012). Although the microtubule cytoskeleton appears as a pivotal factor contributing to neuronal susceptibility in both brain ageing and neurodegenerative conditions, what triggers changes in microtubules during ageing, especially within neurons and their axons, remains elusive.

One further cellular system that naturally declines with age across various organisms, including humans, is the efficiency of the autophagic pathway (Barbosa et al., 2018). Autophagy is a degradation pathway which plays an important role in neuronal homeostasis and maintenance by eliminating and recycling damaged cellular constituents. Prolonged metabolic demands in neurons lead to accumulation of faulty and potentially noxious cellular components that can lead to harmful levels of molecules such as reactive oxygen species (ROS) (De Gaetano et al., 2021; Liu et al., 2017; Maday & Holzbaur, 2016). To deal with this and mitigate damage, neurons heavily rely on basal autophagy. For example, Atg7 is critical for the formation and expansion of autophagic membranes and its loss in mouse sciatic neurons leads to a reduced axonal diameter and the appearance of dystrophic axons without any effect on neuronal death (Kim et al., 2024). Similarly, loss of murine JIP3, which interferes with autophagy by hindering the transport of autophagosomes and lysosomes, also leads to axonal atrophy by inducing the formation of axonal swellings, and the deterioration of the microtubule cytoskeleton of cultured rodent neurons (Cason & Holzbaur, 2023; Rafiq et al., 2022). The above studies suggest that efficient autophagy is particularly important for the maintenance of axonal integrity as we progress through ageing. Reports suggest that autophagy may be inefficient in the human ageing brain, as the expression of core autophagy genes are downregulated with age (Lipinski et al., 2010; Pan et al., 2026). While the age-related decline in autophagy is well-established, it remains unclear how inefficient autophagy can contribute to the remodelling of neuronal networks during physiological ageing, and the specific mechanisms involved in this process.

In this study, we investigate the relationship between autophagy and axonal microtubule deterioration as well as their concerted role in axonal atrophy during ageing. Utilising the *Drosophila* visual system (Okenve-Ramos et al., 2024; Shields et al., 2025), we establish that there is a natural decay of autophagic processes during ageing and that loss of core autophagic factors negatively impact on the microtubule cytoskeleton and axonal maintenance while, enhancing autophagy prevent their natural deterioration during ageing. At the mechanistic level, microtubule defects induced by decreased autophagy, correlates with changes in microtubule bound EB1 protein and can be rescued with application of antioxidants. Enhancing EB1 function was able to rescue autophagy-induced microtubule deterioration and prevents age-related axonal and synaptic atrophy. Our data establishes a cause-effect relationship between autophagy deficiency, microtubule dysregulation and axonal atrophy during ageing. This sheds light into the chain of events that could contribute to neuronal vulnerability during ageing.

## Results

### Autophagy decreases in T1 medulla neurons during ageing

Previously, we showed that T1 medulla neurons in the *Drosophila* brain display typical hallmarks of ageing and comprise morphological changes in axons and terminals (Okenve-Ramos et al., 2024; Shields et al., 2025). Here, we investigated whether autophagy may be deficient in ageing T1 neurons providing a potential mechanism leading to the observed neuronal decay during ageing.

We started by comparing the number and size of autophagic vesicles (AVs) from young and old fly brains. In neurons, autophagosomes predominantly form at distal axonal terminals and mature into degradative organelles by fusing with lysosomes both during their retrograde transport and upon reaching the cell body (Karpova et al., 2025; Maday et al., 2012). To study AVs in terminals and neuronal cell bodies (somas), we targeted the mCherry-tagged, autophagosome marker Autophagy-related 8a (Atg8a; orthologue of mammalian LC3 protein family), using the *GMR31F10-Gal4* driver which expresses in T1 neurons of the optic lobe; (Fig. 1A; (Okenve-Ramos et al., 2024; Qu et al., 2019; Shields et al., 2025)). Atg8a::mCherry labels all AVs including phagophores, autophagosomes and autolysosomes (Nagy et al., 2015). To achieve limited and equal duration of Atg8a-mCherry expression in young and old specimens, we combined *GMR31F10-Gal4* with the conditional expression system Gal4/Gal80^ts^ (Pfeiffer et al., 2010; Suster et al., 2004). Gal80^ts^ is a conditional transcriptional repressor of Gal4, inhibiting its expression at 18°C but allowing expression at 29°C.

**Figure 1.**
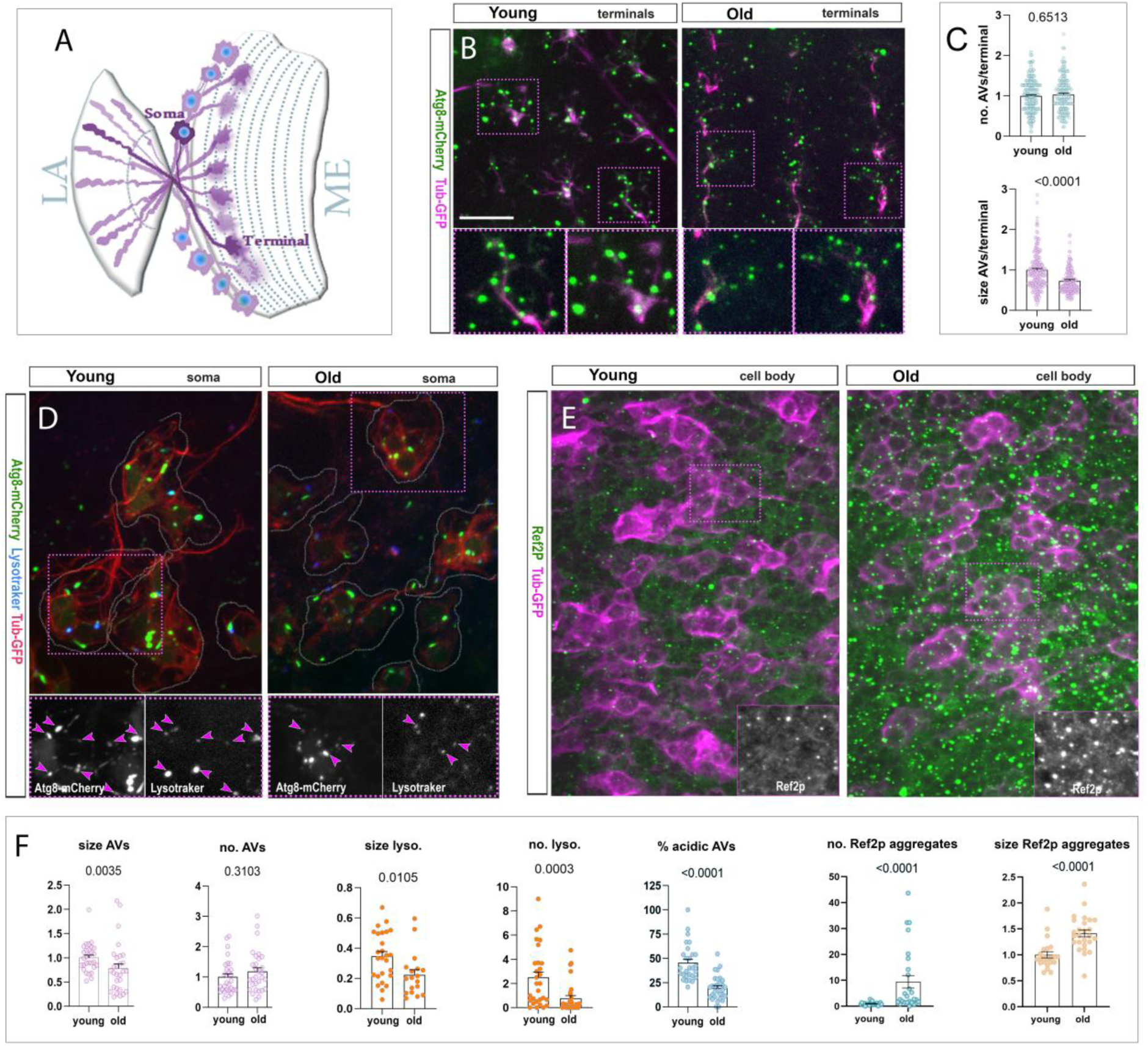
Ageing compromises autophagy in *Drosophila* visual neurons. (**A**) Illustration of, an optic lobe of the adult *Drosophila* brain depicting eight T1 medulla neurons. The T1 somas (S) are located in the medulla (ME), each cell body/soma has a projection that bifurcates terminating in a synaptic terminal in the medulla (terminals) and in the lamina (LA). (**B**) Terminals of the T1 medulla projections from young and old specimens labelled with mCherry tagged Atg8a (Atg8a-mCherry, green) to label autophagic vesicles (AVs) and GFP tagged α-tubulin (Tub-GFP in magenta) for T1 identification. An equal pulse of 6-day expression in old and young specimens is achieved using the UAS/Gal4/Gal80^ts^ system. Expression is silenced at 18⁰C throughout development and adult life until the last 6 days before imaging, at which point flies were shifted to 29⁰C to induce gene expression (expression for 6 days). Young specimens (8-10 day old flies with 2-4 days at 18⁰C “Off” + 6 days at 29⁰C “On”) are compared to old specimens (61-64 days old flies with 55-58-days at 18⁰C “Off”+ 6 days at 29⁰C “On”). (**C**) Quantifications of the number and size (in μm) of terminal AVs from genotypes shown in B. Graphs indicate mean ± SEM, with data points shown as circles; p-values obtained via Mann-Whitney test are indicated in the graphs. Data were taken from a minimum of 12 medullas per group. (**D**) Somas of T1 neurons in the medulla of young and old specimens, expressing with the UAS/Gal4/Gal80^ts^ system, tub-GFP (red), Atg8a-mCherry (green) and labelled with lysotracker (blue). Atg8a-mCherry positive vesicles colocalised with lysotracker (magenta arrowheads in insets), indicate autophagolysosomes (acidic AVs). Young specimens (7-12 day old flies with 1-4 days at 18⁰C “Off” + 6 days at 29⁰C “On”); old flies (49-51 days with 43-45-days at 18⁰C “Off”+ 6 days at 29⁰C “On”). **(E)** Medullas from young (3-6 days at 29°C) and old specimens (30-33 days at 29°C), expressing Tub-GFP in T1 neurons (magenta) and stained with anti-Ref2p antibody (green) depicting Ref2p-positive aggregates. (**F**) Quantifications of the number and size (in μm) of somatic AVs, lysosomes and Ref2p-positive aggregates from genotypes shown in D and E; and of the % of acidic AVs. Graphs indicate mean ± SEM, with data points shown as circles; p-values obtained via Mann-Whitney test are indicated above. Data were taken from a minimum of 7 brains per group. Scale bar in B represents 10 µm in B and E, and 20 µm in D and insets.

For our experimentation, young (6-10 days old at 18°C) and old (64-69 days old at 18°C) flies were reared for 6 days at 29°C to permit expression of Atg8a-mCherry before dissection. Dissected brains were visualised via an *ex vivo* live imaging method (see Materials and Methods) and revealed that AVs were efficiently labelled, both in T1 cell bodies and axonal terminals (Fig. 1B). Comparative analysis of AVs in the T1 neurons’ axonal terminals revealed a decrease in their size in aged brains compared to young brains, whereas their number remained unchanged. Similarly, AVs in the cell bodies of aged brains were also smaller than the young controls (Fig. 1B, C, D and F).

Changes in AV shape might indicate defective autophagy (Jin & Klionsky, 2014). As a direct readout for autophagic activity, we co-labelled Atg8a-mCherry positive AVs with lysotracker. Lysotracker labels acidified vesicles and the presence of both markers in the same vesicle indicates an AV that has fused with a lysosome (autolysosome), as is typical of the late stages of autophagy (Lőrincz et al., 2017). In T1 neurons of aged specimens, we found a pronounced decrease in the percentage of AVs co-labelled with lysotracker, suggesting defects in the acidification of AVs. Furthermore, there is a decrease in the number and size of lysotracker positive vesicles in aged brains (Fig. 1D and F). These results suggest autophagy to be aberrant in aged neurons.

One function e is the removal of intracellular protein aggregates. By immunostaining for Ref(2)P/p62, we observed a substantial increase in the number and size of Ref(2)P/p62-positive puncta throughout the medullas of brains when comparing old to young specimens (Fig. 1E and F). This indicates an accumulation of protein aggregates and supports our notion that basal autophagy is impaired in aged *Drosophila* neurons (Nezis et al., 2008).

### Loss of autophagy regulators induces axonal and synaptic atrophy in T1 neurons

We next asked whether aberrant autophagy might be a driver of axonal and synaptic decay in ageing T1 neurons. First, we tested whether knock-down of autophagy regulators in T1 neurons might exacerbate ageing phenotypes. For this, we used *GMR31F10-Gal4* to express previously validated RNAi constructs against Autophagy-related 1 (*Atg1*) involved in the initiation of autophagy by regulating the formation of autophagosomes to sequester cellular components for degradation (Scott et al., 2007), Autophagy-related 2 (*Atg2*) mediating autophagosomal membrane expansion (Osawa & Noda, 2019) and *Atg8a* involved in various stages of autophagy including membrane tethering, cargo recognition, and autophagosome-lysosome fusion (Mizushima, 2020).

RNAi constructs were co-expressed with the fluorescently tagged membrane marker myristoylated-Tomato (myr-Tom) and axons visualised via *ex vivo* live imaging, to assess for typical ageing hallmarks conserved between vertebrates and *Drosophila*, such as axonal swellings, axonal thinning, and fragmentation of their synaptic terminals (Borzuola et al., 2020; Ceballos-Baumann et al., 1999; Fiala et al., 2007; Okenve-Ramos et al., 2024; Samuel et al., 2011; Shields et al., 2025; Verdú et al., 2000).

When compared to age-matched controls, knock-down of all three autophagy regulators led to T1 axons becoming thinner, developing more axonal swellings and displaying more fragmented and swollen synaptic terminals (Figs. 2A-C and Fig. 1S), thus causing an increase in phenotypes typical of neurons in aged brains. This result, together with the natural decline in autophagy in aged brains, suggest that defective autophagy may be mediating the atrophy of axons and synapses during ageing.

**Figure 2.**
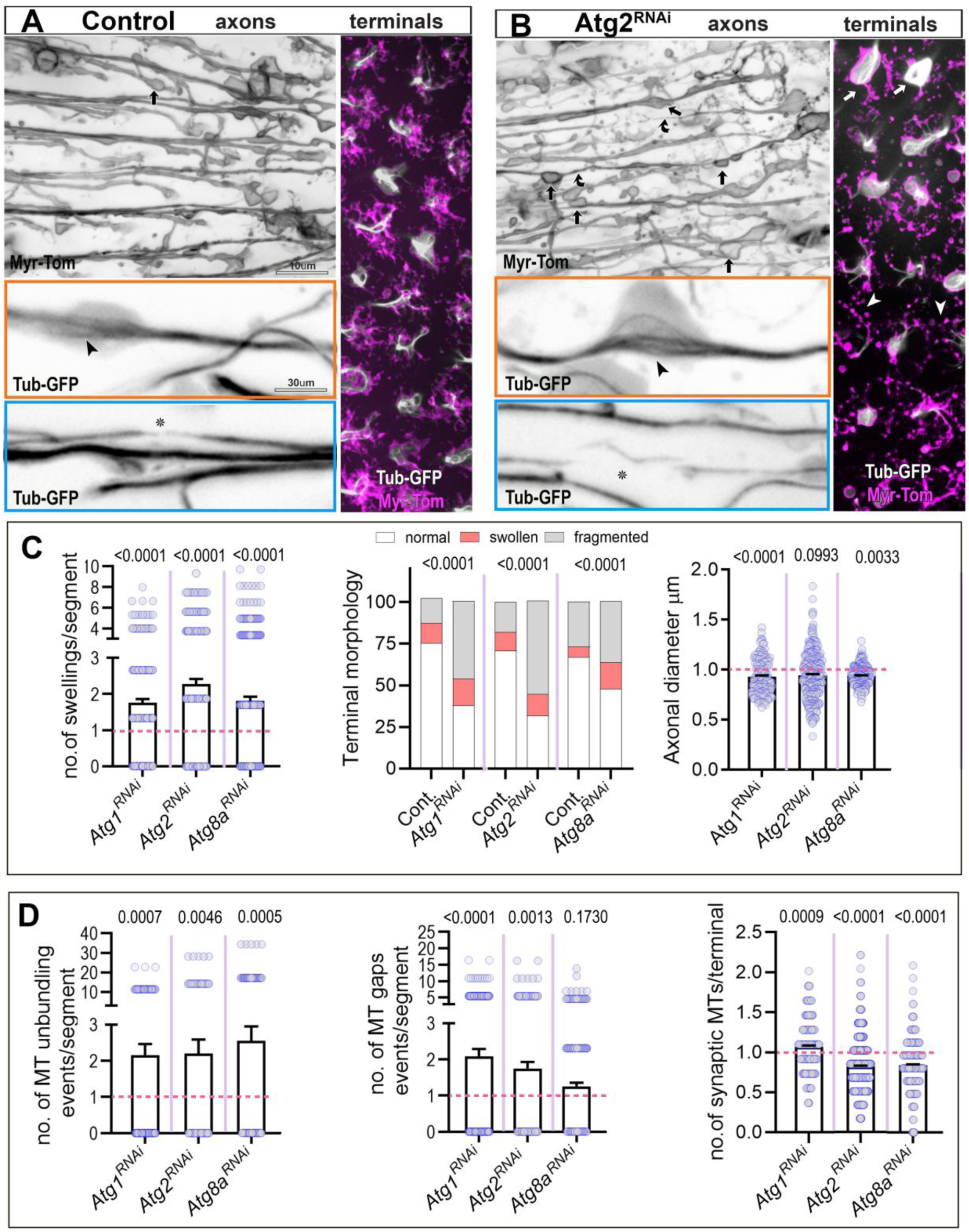
Knocking down Atg1, Atg2 or Atg8a enhances age-related MT deterioration and axonal and synaptic decay in T1 neurons. **(A and B)** Images of T1 axons and terminals in the medulla of 30-33 days old flies labelled with the plasma membrane marker Myr-Tom and GFP-tagged α-tubulin in the absence (A) or presence of Atg2 knockdown (*Atg2^RNAi^* in B). Atg2 knockdown enhances phenotypes typical of ageing neurons, comprising defects in axons such as swellings (black arrows), axon thinning (black bent arrows), MT unbundling (black arrow heads in orange boxes) and gaps in MT bundles (asterisks in blue box); and defects in synaptic terminals including swelling (white arrows) and their breakdown (white arrowhead). (**C and D**) Quantifications of phenotypes shown in A and B plus conditions of further knockdowns for Atg1 (*Atg1^RNAi^*) and Atg8 (*Atg8^RNAi^*). Each knockdown is normalised to its respective control (magenta dashed line). For all measurements except terminal morphology, data points are shown as circles and as mean ± SEM; p-values obtained via Mann-Whitney tests are indicated. For terminal morphology, data are represented as distribution of normal versus swollen or fragmented synapses and significance is obtained via Chi-square test. Data were taken from a minimum of 24 medullas except for axonal diameter where a minimum of 15 medullas were analysed. Scale bars in A represents 10 µm in all images except for 30 µm in coloured boxes.

### Loss of autophagy regulators leads to microtubule cytoskeleton defects in T1 neurons

In aged T1 neurons, the uniform microtubule bundles along axons become disorganised and thinner, and synaptic microtubules become sparse (Okenve-Ramos et al., 2024; Shields et al., 2025), mirroring similar observations in aged primate brains (Cash et al., 2003; Fiala et al., 2007). We previously showed that this microtubule decay is a key driver for the remodelling of neurons during ageing, including axonal thinning, formation of swellings and synaptic atrophy (Okenve-Ramos et al., 2024). We therefore asked whether loss of autophagy regulators affects the microtubule cytoskeleton of T1 neurons, such a finding would suggest that microtubule deterioration in ageing occurs downstream of autophagosomal dysfunction, linking it to the observed axonal decay.

For this, we co-expressed the Atg1, Atg2 and Atg8 knock-down constructs with GFP-tagged α-tubulin (tubulin::GFP; (Grieder et al., 2000)) using the *GMR31F10-Gal4* line. Indeed, all knock-downs caused an increase in foci where axonal microtubules lose their tight, parallel bundle arrangement (from now on referred to as ‘microtubule unbundling’). Furthermore, microtubule bundles became thinner and appeared more often interrupted (referred to as ‘microtubule gaps’; for details see Fig. 2D). At synaptic terminals of axons, microtubules take on a splayed organisation, and the number of splayed microtubules was reduced upon Atg2 and Atg8 knockdowns (Fig. 2A-B and D and Fig. 1S).

These results could have significant implications for the mechanisms of ageing, suggesting that autophagy defects that naturally arise during physiological ageing contribute to the deterioration of the microtubule cytoskeleton and subsequent axonal and synaptic atrophy.

### Increased autophagy ameliorates microtubule decay and axonal atrophy during ageing

Our findings so far suggest that inefficient autophagy is a major cause for axonal deterioration during ageing. We therefore reasoned that, vice versa, we may be able to improve age-related axonal atrophy by pharmacologically or genetically boosting autophagy during ageing.

To test this, we dietarily administered 200 μM of the autophagy-enhancing drug Rapamycin (Bjedov et al., 2010; Juricic et al., 2022). Upon feeding Rapamycin to adult flies, we observed a marked improvement in the age-associated phenotypes of T1 neurons; compared to neurons of age-matched controls, they displayed increased axon calibres, a reduction in axonal swellings, and less fragmented synaptic terminal morphology (Fig. 3A-D and F). Furthermore, microtubule bundles showed reduced gaps and disorganisation and numbers of splayed microtubules in synaptic terminals were increased (Fig. 3A-F).

**Figure 3.**
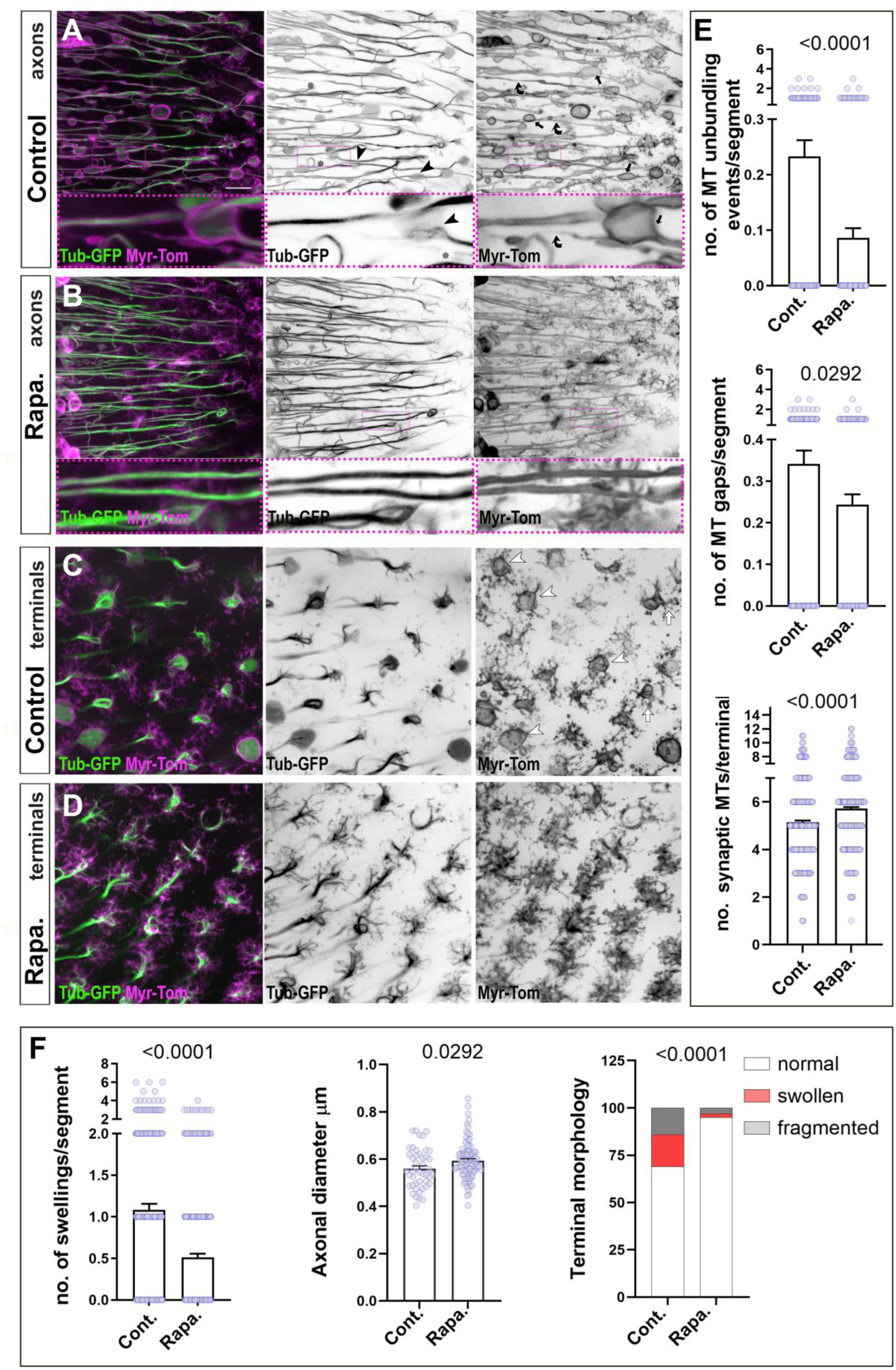
Rapamycin ameliorates axonal and synaptic ageing phenotypes. (**A-D**) T1 axons (A and B) and terminals (C and D) in the medulla of 30-32 days old specimens, labelled with Myr-Tom and Tub-GFP. Ageing phenotypes including axon swellings (black arrows), axon thinning (curved bent arrows), MT unbundling (black arrow heads), gaps in MT bundles (asterisks) and defects in synaptic terminals (including swelling: white arrows and breakdown: white arrowhead) can be observed in old specimens in A (axons) and C (terminals), but are decreased upon dietary administration of 200 mM Rapamycin (B, axons and D, terminals). (**F-E**) Quantifications of ageing phenotypes. For all measurements except terminal morphology, data points are shown as circles and as mean ± SEM; p-values obtained via Mann-Whitney tests are indicated. For terminal morphology, data are represented as distribution of normal versus swollen or fragmented synapses with significance obtained via Chi-square test. Data were taken from a minimum of 27 medullas except for axonal diameter where a minimum of 17 medullas were analysed. Scale bar in A represents 10 µm in all images except for 30 µm in coloured boxes.

Rapamycin acts by inhibiting the mTOR signalling pathway, which is known to suppress autophagy, but also play important roles in other non-autophagy-related processes (Li et al., 2014). To pinpoint autophagy more specifically, we next utilised Atg8a overexpression as a direct way to enhance autophagic activity (Simonsen et al., 2008). Continuous overexpression of Atg8a in T1 neurons likewise improved all the ageing hallmarks listed above and protected microtubules from decay during ageing (Fig. 4A-E), thus corroborating the findings with Rapamycin (Fig. 4A-D and F).

**Figure 4.**
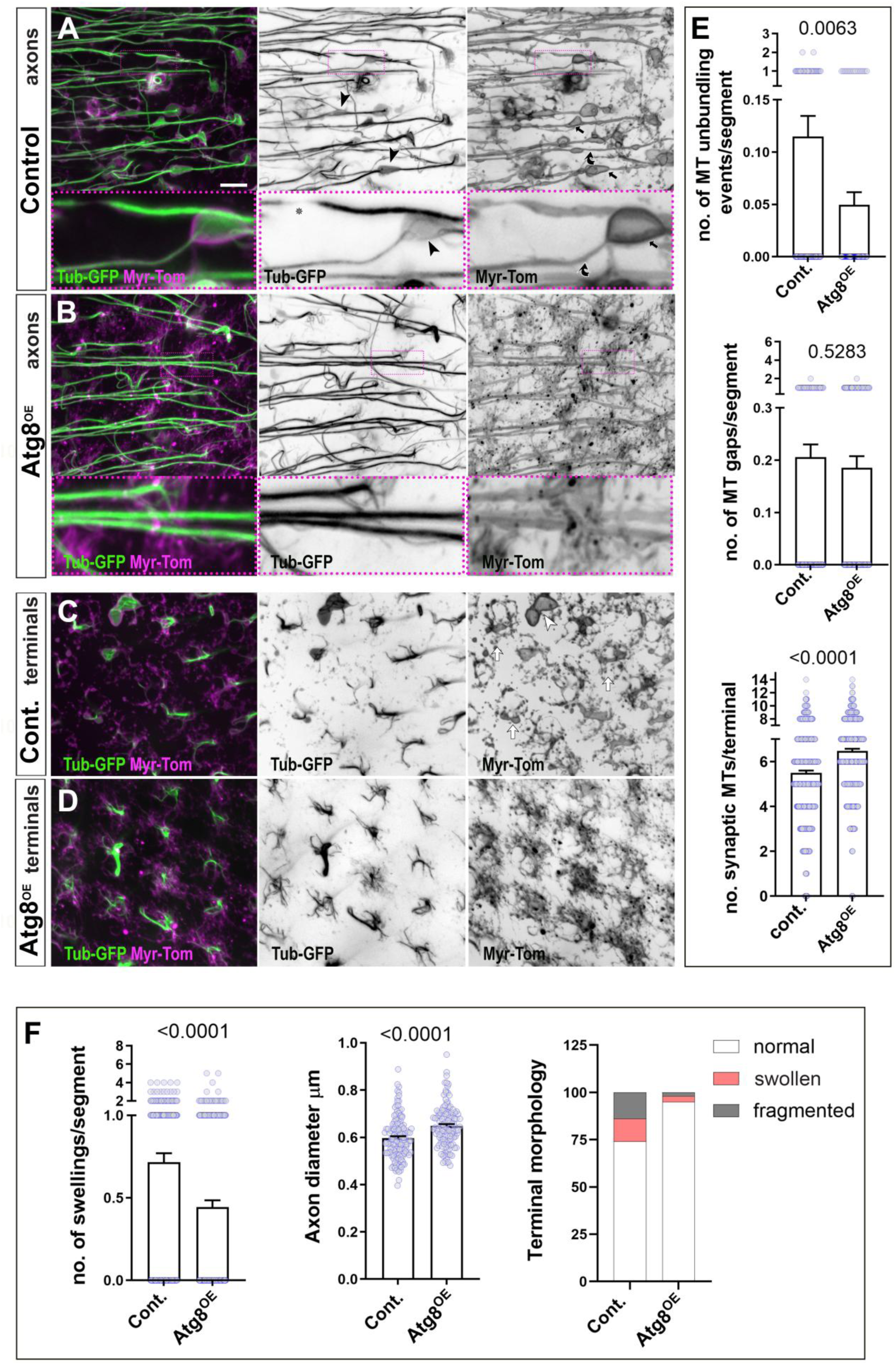
Over-expression of Atg8 improves axonal and synaptic deterioration. T1 axons (A and B) and terminals (C and D) in the medulla of old 31-36 days old fly brains, labelled with Myr-Tom and Tub-GFP. Ageing phenotypes including axon swellings (black arrows), axon thinning (black bent arrows), MT unbundling (black arrow heads) and defects in synaptic terminals (including swelling: white arrows and breakdown: white arrowhead) can be observed in old specimens in A and C but are diminished upon T1 specific over-expression of Atg8a (B,D). (**F and E**) Quantifications of ageing phenotypes. For all measurements except terminal morphology, data points are shown as circles and as mean ± SEM; p-values obtained via Mann-Whitney tests are indicated. For terminal morphology, data are represented as distribution of normal versus swollen or fragmented synapses with significance obtained via Chi-square test. Data were taken from a minimum of 29 medullas except for axonal diameter where a minimum of 20 medullas were analysed. Scale bar in A represents 10 µm in all images except for 30 µm in coloured boxes.

Our findings that upregulation of autophagy can reduce the appearance of all ageing hallmarks assessed, further supports the view that decrease in autophagy is a major cause for axonal atrophy during ageing and that the deterioration of microtubule bundles may be involved in this process.

### Microtubule disorganisation induced by dysfunctional autophagy is rescued by the antioxidant Trolox in cultured neurons

Having identified microtubule bundle decay as a potential mediator linking dysfunctional autophagy to axonal atrophy, we aimed to explore potential mechanisms involved, capitalising on *Drosophila* cultured primary neurons as an efficient and well-established experimental model (Prokop et al., 2012; Sánchez-Soriano et al., 2010; Voelzmann & Sanchez-Soriano, 2022).

We first treated cultured neurons with Chloroquine and Bafilomycin A1, two drugs known to block autophagy by inhibiting autophagosome-lysosome fusion and preventing the activation of pH-sensitive lysosomal hydrolases (Mauvezin & Neufeld, 2015; Redmann et al., 2017). We analysed primary neurons fixed at 6 days in vitro (DIV) that were exposed to 100 μM Chloroquine for the last 3 days of the culture period, or neurons fixed at 1 DIV treated with 100 nM Bafilomycin throughout the culture period. When compared to vehicle-treated controls, the axons of neurons treated with Chloroquine or Bafilomycin A1 displayed prominent microtubule curling and unbundling (Fig. 5; quantified as microtubule disorganisation index, MDI; see Methods). The same microtubule disorganisation phenotypes were observed in neurons cultured from embryos homozygous for either the *Atg8a^KG07569^* or the *Atg1^Δ3d^* loss-of-function mutant alleles (Kołodziej et al., 2024; Pimenta de Castro et al., 2012; Scott et al., 2007; Tan et al., 2024) (Fig. 5 and 6). These data jointly suggest that decreased autophagy function was sufficient to interfere with microtubule regulation *in vitro* thus reproducing our findings with T1 neurons in the adult fly brain, which displayed very similar microtubule disorganisation upon ageing or knock-down of autophagy-linked genes (Fig. 2-4).

**Figure 5.**
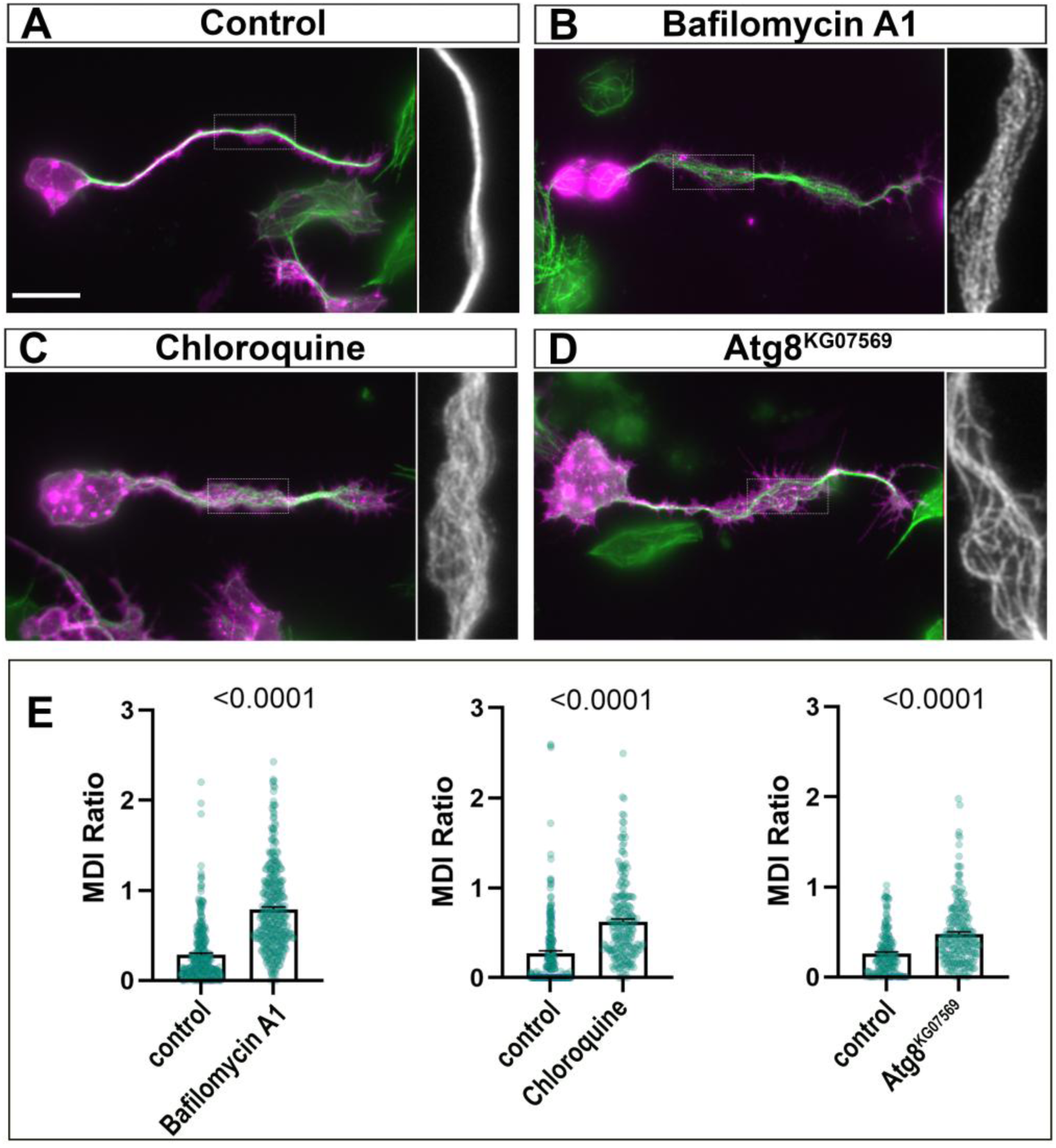
Defective autophagy induces microtubule disorganisation in primary neuronal cultures. **(A-D)** Representative images of *Drosophila* primary neurons stained for tubulin (green) and Hrp (magenta). Neurons at different conditions: control **(A)**, control treated with 100 nM Bafilomycin A1 **(B)**, control treated with 100 μM Chloroquine **(C)**, and neurons carrying the loss of function mutant allele *Atg8a^KG07569^* in homozygous **(D)**. Stippled boxes are shown as magnified single MT channel and rotated 90 degrees clockwise. **(E)** Quantitative analysis of the microtubule disorganisation index (MDI). Each drug treatment is compared to controls exposed to the drug vehicle (DMSO for Bafilomycin A1 and water for Chloroquine). Bars represent normalised mean ± SEM. p-values are shown above each bar, as assessed using Mann-Whitney tests. Data were collated across a minimum of 2 experimental repeats with 3 individual cultures per condition per repeat, with a minimum of 180 neurons evaluated per condition. Scale bars = 5 μm.

**Figure 6.**
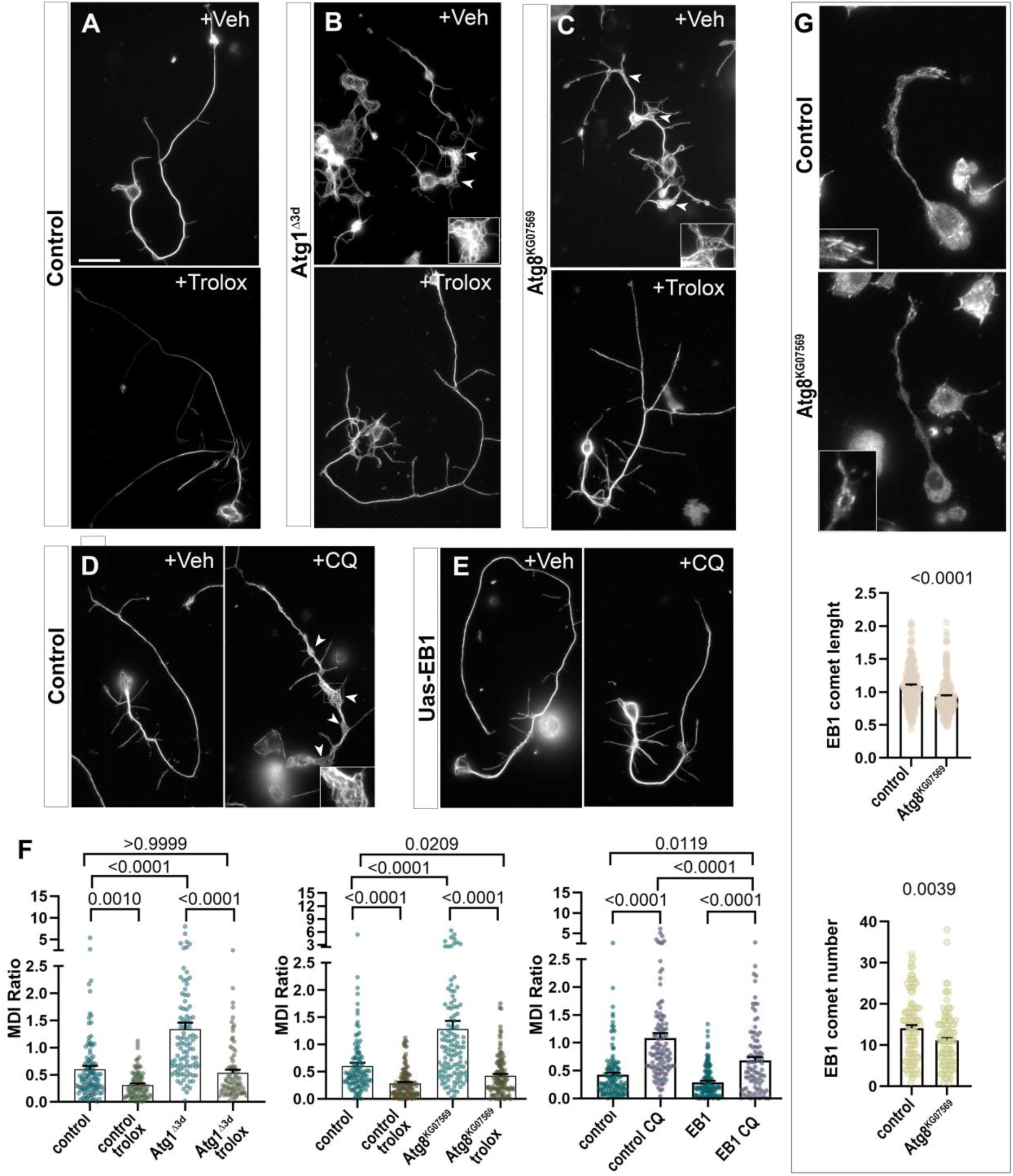
Microtubule disorganisation caused by autophagy loss is rescued by the antioxidant Trolox or by overexpression of EB1 in primary neuronal cultures. (A-E) Representative images of *Drosophila* primary neurons 6 DIV stained for tubulin. (A-C) Control neurons and neurons carrying the mutant allele *Atg1^Δ3D^* or *Atg8a^KG07569^* in homozygous are treated with 100 μM Trolox or Ethanol (vehicle, Veh). (D-E) Control neurons and neurons overexpressing EB1 with the Elav-gal4 driver, are treated with 100 μM Chloroquine or vehicle (water). White arrow heads indicate regions of microtubule disorganisation, and dashed boxes magnified insets. (F) Quantitative analysis of microtubule disorganisation with the MDI. Bars represent normalised mean ± SEM. p-values are shown above each bar, as assessed by a Kruskal-Wallis one-way test. (G) *Drosophila* primary neurons at 6 HIV (hours *in vitro*) stained for EB1. Control neurons and *Atg8a^KG07569^* neurons, with EB1 comets magnified in the respective inset and quantitative analysis of EB1 comet lengths (in μm) per cell and number per axon; bars represent mean ± SEM normalised to control; p-values are shown above each bar, as assessed by Mann-Whitney tests. Data were collated across a minimum of 2 experimental repeats with 3 individual cultures per condition per repeat, with a minimum of 100 neurons evaluated per condition. Scale bars = 10 μm.

Microtubule disorganisation can be triggered by the dyshomeostasis of reactive oxygen species (ROS) (Liew et al., 2026b; Murray-Cors et al., 2025; Shields et al., 2025), and dysfunctional autophagy is a known cause for oxidative stress (Chang et al., 2022; Filomeni et al., 2015; Gouda et al., 2025; Park et al., 2021; Sedlackova et al., 2020; Yan & Finkel, 2017; Zhao et al., 2019). We therefore investigated whether microtubule disorganisation downstream of defective autophagy might be triggered by ROS dyshomeostasis. For this, we treated *Atg8a^KG07569^* and *Atg1^Δ3d^* homozygous mutant neurons for 3 days with 100 μM of the vitamin E analogue Trolox, which is an effective ROS scavenger, hence antioxidant (Brigelius-Flohé & Traber, 1999; Giordano et al., 2020). When analysed at 3 DIV, vehicle-treated *Atg1^Δ3d^* or *Atg8a^KG07569^* mutant neurons showed significant microtubule disorganisation phenotype, but this was suppressed in the mutant neuron cultures treated in parallel with Trolox (Fig. 6 A-C and F).

Taken together, these data suggest that defective autophagy disrupts the organisation of axonal microtubules and this is mediated by excessive ROS.

### The microtubule regulator Eb1 protects ageing neurons from the detrimental effects of defective autophagy

We previously showed that increased ROS interferes with the function of End-binding protein 1 (Eb1) (Shields et al., 2025). Eb1 associates with plus ends of polymerising microtubules (often referred to as Eb1 comets), regulating their dynamics and stability (Duellberg et al., 2016; Goodson & Jonasson, 2018; Rickman et al., 2017). Functional deficiency of Eb1 causes microtubule disorganisation in *Drosophila* primary neurons (Alves-Silva et al., 2012; Hahn et al., 2021). We therefore hypothesised that the impact of defective autophagy on microtubules could involve Eb1 dysfunction. To test this, we measured the intensity of endogenous Eb1 comets at microtubule plus ends in cultured neurons. Compared to controls, the length and number of Eb1 comets was significantly reduced in neurons homozygous for *Atg8a^KG07569^* (Fig. 6G).

We next established whether Eb1 overexpression might protect from microtubule disorganisation under conditions of decreased autophagy. For this, cultured neurons with and without targeted Eb1 overexpression were treated with 100 μM Chloroquine (administered as explained above); we found that Eb1-overexpressing neurons displayed a significant decrease in Chloroquine-induced microtubule disorganisation (Fig. 6E and F), consistent with a protective effect of Eb1.

Following the protective effects of EB1 observed *in vitro*, we investigated whether Eb1 could similarly rescue autophagy-defective neurons *in vivo.* Since Chloroquine administered through feeding, has been shown to inhibit autophagy also in adult flies (Bargiela et al., 2019; Nagy et al., 2018; Zirin et al., 2013), we used this approach to test protective effects of Eb1 overexpression also in T1 neurons.

As a first step, we dietarily administered Chloroquine to adult flies during ageing. These flies displayed a significant increase in ageing hallmarks when compared to age-matched controls, and this included increase in axonal swellings and further deterioration of synaptic terminals. We also observed that Chloroquine treatment significantly increased the deterioration of microtubules in axons and synaptic terminals (details in Fig. 7). In contrast, Chloroquine-fed flies with targeted Eb1 overexpression in T1 neurons displayed a significant reduction in all these ageing hallmarks, including their microtubules, axonal swellings and synaptic morphology (Fig. 7).

**Figure 7.**
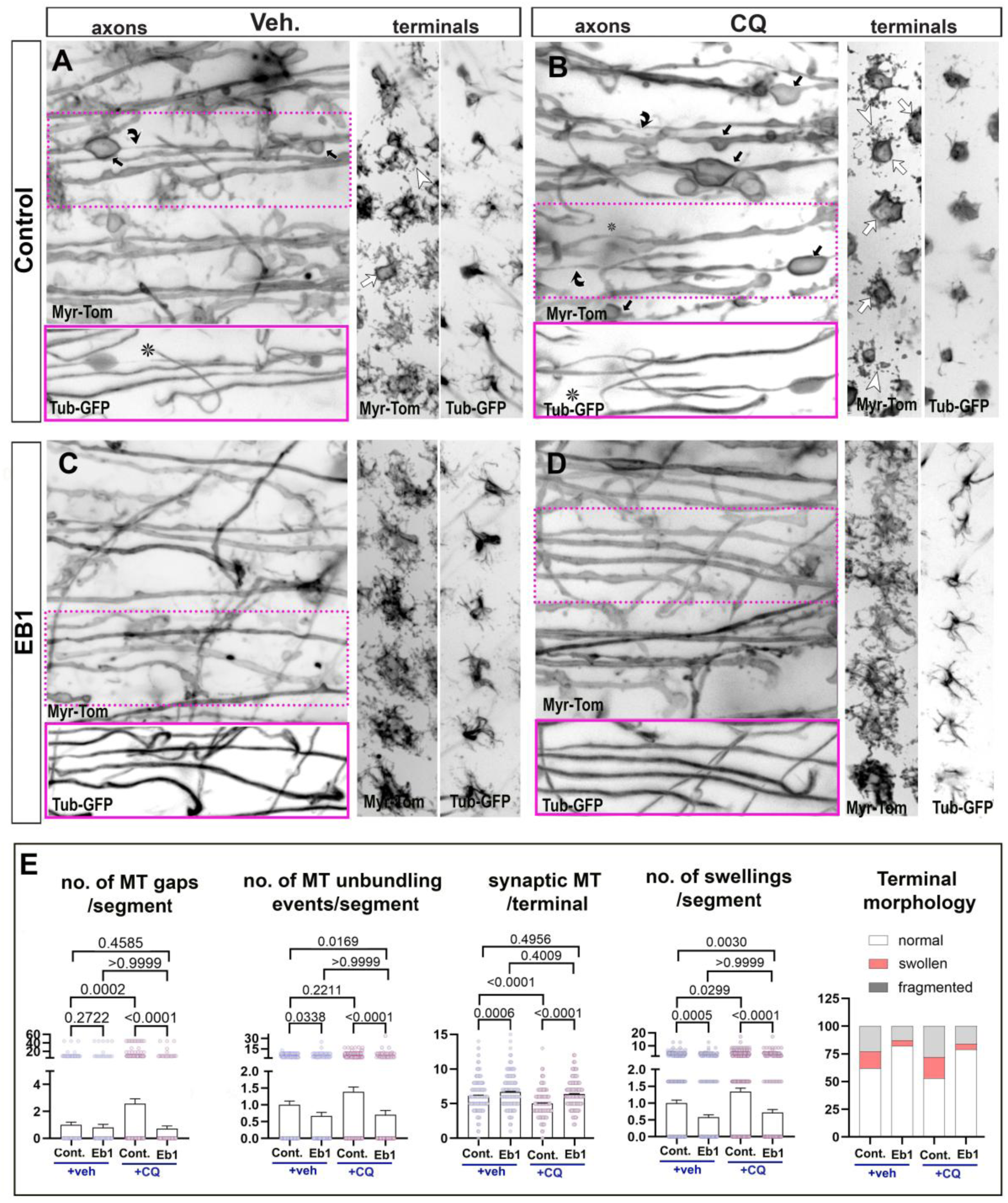
EB1 overexpression ameliorates CQ-induced axonal and synaptic ageing phenotypes. **(A-D)** T1 axons and terminals in the medulla of 27-29 days old specimens, labelled with Myr-Tom and Tub-GFP. **(A and B)** Ageing phenotypes including axon swellings (black arrows), thinning of axonal MT bundles (asterisks), synaptic deterioration (swollen and fragmented morphologies, white arrowheads and arrows, respectively) and reduced synaptic microtubules, are exacerbated in old specimens exposed to dietary addition of 100 μM CQ (exposure for 25-27 days). **(C and D)** T1 specific overexpression of EB1 protects neurons from CQ enhancement of ageing phenotypes. (**E**) Quantifications of ageing phenotypes. For all measurements except terminal morphology, data points are shown as circles and as mean ± SEM; p-values obtained via Kruskal-Wallis one-way test. For terminal morphology, data are represented as distribution of normal versus swollen or fragmented synapses with significance obtained via Chi-square test. Data were taken from a minimum of 26 medullas. Scale bar in A represents 10 µm in all images except for 30 µm in coloured boxes.

Taken together, we established a mechanistic cascade that leads from autophagy dysfunction to ROS dyshomeostasis which, in turn, inhibits Eb1 thus causing microtubule deterioration that, eventually, leads to axonal atrophy. In this cascade, we show that promoting autophagy or stabilising microtubules via Eb1 overexpression can interrupt the detrimental chain and prevent or slow down the occurrence of ageing hallmarks.

## Discussion

In this study, we demonstrate that ageing of medulla neurons in the *Drosophila* brain is characterised by altered autophagic vesicle morphology, reduced lysosomal acidification, and accumulation of Ref(2)P/p62 aggregates, collectively indicating a failure of cellular clearance and protein homeostasis. Crucially, we identified a functional dependency between defective autophagy and neuronal decay during physiological ageing. Our detailed further analyses of this phenomenon revealed a mechanistic cascade where defective autophagy causes microtubule decay which, in turn, leads to axon atrophy, as reflected in axon thinning, appearance of swellings and synaptic decay. Furthermore, we show that preserving the microtubule cytoskeleton mitigates the deleterious effects of declining autophagy.

Our data suggest key features of this mechanistic cascade where defective autophagy causes ROS dyshomeostasis and, in agreement with our previous work (Shields et al., 2025), inhibits Eb1 as a key regulator of the microtubule cytoskeleton in axons (Alves-Silva et al., 2012; Hahn et al., 2021). Consequently, the microtubule cytoskeleton emerges as a key target of age-related autophagic failure.

Taken together, our work integrates these disparate features of ageing (defective autophagy, increase in ROS and microtubule deterioration) into a causative chain of events. The finding that strengthening any step in this pathway diminishes the hallmarks of ageing, achieved through interventions in this study such as enhancing autophagy or expressing Eb1 (known to promote microtubule bundling during ageing (Okenve-Ramos et al., 2024)), or antioxidant treatments (Shields et al., 2025), suggests that brain decay during ageing is a preventable process.

### Microtubules as a key downstream mediator from deficient autophagy to axon atrophy

Structural and morphological alterations in neurons, particularly within dendrites and axons, are significant features of the aged brain that drive the gradual decay of neuronal function (Dickstein et al., 2013; Morrison & Baxter, 2012). Our study underscores the fundamental role of autophagy in driving this process. By establishing a causative link between autophagic dysfunction and age-related neuronal atrophy, our findings support the growing consensus that autophagic deficiency is a defining hallmark of the aged brain. This is consistent with evidence that autophagic activity naturally declines during ageing across species, from invertebrates to humans (Aman et al., 2021; Karpova et al., 2025; López-Otín et al., 2023; Pan et al., 2026; Schmid et al., 2024; Stavoe et al., 2019; Yang et al., 2023).

Our detailed cell biology analysis demonstrates a significant decrease in autophagic activity in the *Drosophila* ageing brain. At axonal synaptic terminals, we observed reduced AV size, suggesting that autophagic initiation and biogenesis may be compromised by the inability of autophagosomes to expand or mature, which is consistent with reports of stalled autophagosome formation in the distal axons of aged neurons (Stavoe et al., 2019; Tsong et al., 2023). However, the most profound alterations occurred within the cell bodies of aged T1 neurons, characterised by a marked reduction in acidified AVs and lysosomes. These observations suggest that the primary defect may be a breakdown in autophagosome maturation and lysosomal fusion. Similar declines in the degradative capacity of somatic lysosomes through deacidification are seen in murine neurons ageing *in vitro* (Burrinha et al., 2023) and in early-stage Alzheimer’s disease models (Lee et al., 2022).

The functional importance of the deterioration of this degradative pathway is highlighted by our finding that inhibiting autophagy (knocking down Atg1, Atg2 or Atg8a, or feeding Chloroquine) accelerates ageing hallmarks. Conversely, promoting autophagy through Atg8a overexpression or Rapamycin prevents axonal and synaptic decay. These results align with the murine studies where Chloroquine exacerbates ageing-like phenotypes at the neuromuscular junction (Gulbronson et al., 2025), while Spermidine or Rapamycin supplementation enhances cognitive performance and mitigates neurodegenerative pathology in various aged models (Cai et al., 2024; Jahrling & Laberge, 2015; Ni & Liu, 2021; Schroeder et al., 2021; Sigrist et al., 2014).

Despite the growing importance of this ageing hallmark, it is not yet clear how autophagy causes neuronal remodelling and atrophy during ageing. Our work helps to bridge this gap by suggesting a mechanistic cascade in which defective autophagy triggers microtubule decay ultimately leading to axonal and synaptic atrophy during ageing. We previously demonstrated that microtubule deterioration is a critical event mediating axonal and synaptic decay during ageing (Okenve-Ramos et al., 2024; Shields et al., 2025); here, we show that proper functioning autophagy is essential to sustain adequate microtubule bundles in ageing neurons. The exacerbation of microtubule deterioration in autophagy-deficient mutants, and their improvement provided by enhanced autophagy in aged brains, identifies insufficient autophagy as a primary driver of this cytoskeletal deterioration. Furthermore, the finding that Eb1 expression, which improves microtubule bundle health, prevents atrophy even in autophagy-deficient conditions, further confirms the microtubule cytoskeleton as a key downstream mediator in this cascade. Given that both reduced autophagy and microtubule deterioration are conserved in the human ageing brain (Aman et al., 2021; Cash et al., 2003; Fiala et al., 2007; López-Otín et al., 2023; Schmid et al., 2024), these findings provide a vital mechanistic microtubule-mediated link between metabolic health and the physical integrity of aged neurons.

### Mechanistic links between deficient autophagy and microtubule decay

We proposed that an increase in ROS, caused by defective autophagy, could act as a pathological mediator between autophagic failure and microtubule deterioration. Under normal conditions, autophagy limits oxidative stress, for example by sequestering damaged mitochondria (mitophagy), peroxisomes (pexophagy) and oxidised protein species (Filomeni et al., 2015; Ornatowski et al., 2020). When autophagy declines with age, these pro-oxidant sources accumulate, leading to a surge in local ROS levels within neurons (Gouda et al., 2025; Tran & Reddy, 2020; Wei et al., 2025) and our previous work has shown that ROS compromises axonal integrity by triggering premature decay of the microtubule cytoskeleton that precedes neuronal atrophy during ageing (Shields et al., 2025). Since microtubule decay upon deficient autophagy is rescued by the antioxidant Trolox, we suggest ROS as a likely intermediary. We propose that when autophagy fails, the resulting accumulation of damaged organelles or proteins triggers an oxidative burst that directly targets microtubule properties.

There is significant literature support linking ROS, microtubule deterioration, and neuronal ageing. Independent studies in vertebrate systems and *in vitro* models confirm that oxidative stress is a direct driver of cytoskeletal instability and axonal decay (Beckhauser et al., 2016; Conze et al., 2025; Drum et al., 2016; Gardiner et al., 2013; Liew et al., 2026a; Murray-Cors et al., 2025; Wilson & González-Billault, 2015). A likely target of ROS in this context is Eb1 which acts as a scaffold that guides extending microtubules into organised bundles; Eb1 deficiency causes microtubule unbundling and disorganisation as a prominent hallmark of axon decay (Alves-Silva et al., 2012; Hahn et al., 2021; López-Otín et al., 2023; Okenve-Ramos et al., 2024). Eb1’s ability to bind and regulate microtubules is compromised by high levels of ROS (Shields et al., 2025) and defective autophagy (Fig. 6b), and overexpressing Eb1 can bypass the detrimental effects of ROS (Shields et al., 2025) and of deficient autophagy during ageing (Fig. 7). How Eb1 function is affected by oxidative stress is currently unknown, but it is likely to involve the post-translational modification or oxidisation of tubulin and/or Eb1, and potentially also other microtubule-associated proteins (MAPs). The Eb1-mediated rescue demonstrates that we can bypass the inhibitory signals generated by aberrant autophagy and oxidative stress.

### Conclusions and future perspectives

Altogether, our work supports the notion that an age-related decline in autophagy acts as a primary driver of brain ageing. While ageing is universally recognised as a critical contributor to late-onset, sporadic neurodegeneration, the exact mechanisms rendering aged neurons vulnerable to stressors have remained elusive. Most significantly, our findings bridge this gap by demonstrating that defective autophagy acts as a direct causal trigger for the loss of microtubule integrity which drives downstream neurite and synapse atrophy. Uncovering this specific cascade offers new therapeutic possibilities. By targeting microtubules as the last link in the pathological chain, interventions could likely integrate and mitigate other ROS and autophagy related pathologies, thereby exerting the widest possible protective effect. Ultimately, these findings position autophagy and microtubule enhancers as viable candidates for maintaining axonal function and synaptic connectivity, not only in the normal ageing brain but also across a broad spectrum of neurodegenerative conditions characterised by impaired autophagy, lysosomal function or disrupted ROS homeostasis.

## Methods

### Fly stocks and husbandry

Flies were maintained on standard fly food containing cornflour, glucose, yeast and agar. For experiments involving ageing, flies were maintained at 25°C during development and transferred to 29°C post-eclosion to expedite the ageing process. Flies were maintained at low densities (maximum of 20 flies per vial) and were transferred to new vials with fresh food every 3 days. Females were used for this study.

The following fly stocks were used in this project. Gal4 driver lines include: *elav-Gal4* (3^rd^ chromosomal, expressing pan-neuronally at all stages; (Luo et al., 1994)), *GMR31F10-Gal4* (Bloomington #49685; expressing in T1 medulla neurons; (Okenve-Ramos et al., 2024; Qu et al., 2019; Shields et al., 2025). Mutant stocks: *Atg1Δ3D* (kind gift from Dr. Samantha Loh) and *Atg8^KG07569^* (Bloomington stock #14639). Lines for transgene targeted expression were *UAS-EB1-GFP* (Alves-Silva et al., 2012), *UAS-GFP-α-tubulin84B* (Grieder et al., 2000), *UAS-*Atg8a-mCherry (Nagy et al., 2015) (Bloomington stock #37750) and *UAS-myr::tdTomato* (Bloomington stock #32222). Lines for targeted gene knockdowns were acquired from Bloomington Drosophila Stock centre including *Atg1 RNAi* (Bloomington stock #26731), *Atg2 RNAi* (Bloomington stock #27706) *and Atg8 RNAi* (Bloomington stock #28989).

### Adult drug feeding

Chloroquine (Sigma Aldrich) feeding protocols were adapted from (Bargiela et al., 2019). After eclosion, 2-day old flies were transferred to vials containing either 100 μM Chloroquine (mixed in standard fly food) or water (vehicle, mixed in standard fly food). Flies were maintained on Chloroquine or water till the day of imaging and tipped on fresh food every 2-3 days. Flies were then prepared for confocal imaging (*described below*). Similar protocol was adapted for feeding either 200 μM Rapamycin (LC laboratories) or the vehicle DMSO mixed in standard fly food.

### Adult brain dissections and ex vivo preparation, microscopy and image analyses

Dissections of *Drosophila* adult brains were performed in Dulbecco’s PBS (Sigma, RNBF2227) after briefly anaesthetising the flies on ice. For live imaging, dissected brains were placed into a drop of Dulbecco’s PBS on MatTek glass bottom dishes (P35G1.5-14C), with a spacer and covered by coverslips. Brains were imaged immediately.

For lysotracker treatment, dissected brains were incubated in the dark for 40 minutes on 100nM lysotracker deep red diluted in Dulbecco’s PBS.

To measure ageing hallmarks in the optic lobe of adult flies, brains were dissected as explained above and immediately imaged with a 3i Marianas Spinning Disk Confocal Microscope at the Centre for Cell Imaging at the University of Liverpool. A section of the medulla columns comprising the 4 most proximal axonal terminals was used to quantify axonal microtubule phenotypes and the number of swellings. The line tool of ImageJ was used to measure the diameter of axons and their MT core, at evenly spaced unbiased positions predetermined by a grid. A mean per axon from multiple measure points was obtained. To score synaptic readouts, a square was drawn through the central part of the medulla and 30-50 synapses were scored per medulla. Synaptic MTs and morphological categorization were calculated manually. For quantifications of AVs and lysotracker positive vesicles, cell bodies and terminals were selected in ImageJ. The 3D mitochondria analyser plugin for ImageJ/Fiji was used to analyse complex 3D organelles using adaptive thresholding to identify vesicular objects. For colocalization studies of Atg8a-mCherry and lysotracker, the DiAna plugin for ImageJ/Fiji was used. For quantifications of Ref(2)P/p62 positive aggregates, an equal size area of each brain was selected, and the average background was subtracted. For this an average background was calculated for each brain from 10 independent readings. Adaptive threshold followed by Analyze particles from ImageJ/Fiji was used to identify Ref(2)P aggregates.

### Primary neuronal cell culture and drug treatments

*Drosophila* primary neuronal cultures were carried out using methods previously described (Prokop et al., 2012; Sánchez-Soriano et al., 2010; Voelzmann & Sanchez-Soriano, 2022). Drug treatments were carried out at 0–3 DIV, as stated throughout according to the experimental approach. Compounds used were: 100 µM Chloroquine (vehicle: water), 100 nM Bafilomycin A1 (vehicle: DMSO), and 100 µM Trolox ([±]-6-Hydroxy-2,5,7,8-tetramethylchromane-2-carboxylic acid; Merck, vehicle: ethanol).

### Immunocytochemistry of brains and cultured neurons

Cultured neurons were fixed in 4% paraformaldehyde (PFA) in 0.05 M phosphate buffer (pH 7.0–7.2) for 30 minutes at room temperature. For analyses of Eb1 comets, primary *Drosophila* neurons were fixed in ice-cold (stored at –80°C) +TIP-fixative solution (90% methanol, 3% formaldehyde, 5 mM sodium carbonate, pH 9; (Hahn et al., 2021; Rogers et al., 2002) for 10 minutes at −20°C. Antibody staining and washes were performed with PBT (PBS supplemented with 0.3% Triton X-100).

*Drosophila* brain dissections were performed as explained above and fixed in 4% paraformaldehyde (PFA; in 0.05 M phosphate buffer, pH 7–7.2) for 30 min at room temperature (RT). Brains were washed in PBT (PBS with 0.3% Triton X-100). Antibody staining and washes were performed with PBT.

Primary antibodies used in this study were: Cy3-751 conjugated anti-HRP (Jackson ImmunoResearch), anti-α-tubulin (mouse, 1:1000; T9026, Merck), anti-Ref2P (rabbit, 1:500; ab178440 from Abcam), anti-DmEB1 (rabbit, 1:2000, gift from H. Ohkura (Elliott et al., 2005)). Secondary antibodies used were FITC-, Cy3- or Cy5-conjugated secondary antibodies (goat, Jackson ImmunoResearch). Cells were mounted in Vectashield (H-1000, VectorLabs).

### Microscopy of cultured neurons and image analyses

Fixed *Drosophila* cells were captured using a Nikon eclipse 90i with a Retiga 3000 camera (QImaging) or a 3i Marianas Spinning Disk using a 63X 1.4 NA Zeiss Plan Apochromat lens and FLASH4 sCMOS (Hamamatsu) camera at the Centre for Cell Imaging at the University of Liverpool. To measure the extent of microtubule disorganisation or curling in axons, a ‘microtubule-disorganisation index’ (MDI) was used. Using FIJI/ImageJ, the areas of disorganised MTs per neuron were marked and measured using the freehand selection tool. The total area of microtubule disorganisation was then divided by the respective axon length, (measured using the segmented line tool) (Prokop et al., 2012; Sánchez-Soriano et al., 2010; Voelzmann & Sanchez-Soriano, 2022). Eb1 comet lengths were measured using the FIJI/ImageJ line tool and data were presented as mean Eb1 comet length per cell. Only axonal Eb1 comets were quantified.

### Statistical analyses

Statistical analyses were performed in GraphPad Prism; statistical tests used are stated in the relevant figure legends. Exact p-values are indicated in graphs. Sample sizes and other statistical values are indicated in figure legends.

## Acknowledgments and disclosures

This work was made possible through support to N.S.S by the BBSRC (BB/R018960/1, BB/W016907/1), to E.G by the NLD BBSRC DTP, BB/T008695/1, and to A.M by the NLD BBSRC DTP3, BB/T008695/1. We thank the University of Liverpool Imaging Facility for assistance during the project. We also thank Sylvie Urbe, Michael Clague, Hiro Ohkura, Dr. Samantha Loh, the fly facility at the University of Cambridge, Gábor Juhász and the fly facility at the University of Manchester for sending us reagents. Stocks obtained from the Bloomington *Drosophila* Stock Center (NIH P40OD018537) were used in this study.

---

**Supplementary figure 1.**
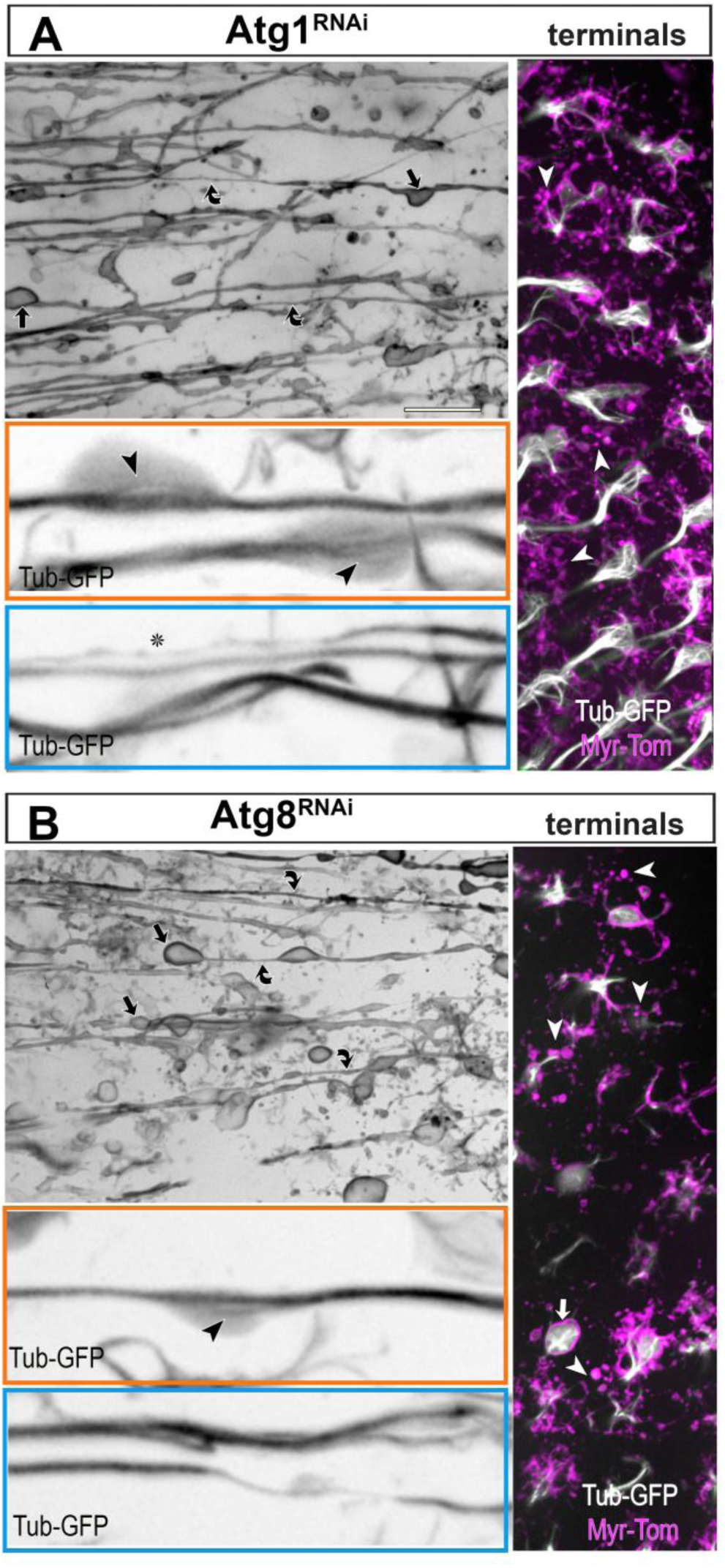
Effect of Atg1 or Atg8a knock down in T1 neurons. **(A and B)** Images of T1 axons and terminals in the medulla of flies expressing *atg1^RNAi^* (35-37 days old) or *atg8^RNAi^* (30-33 days old) together with myr-Tom and GFP-tagged α-tubulin. Both knockdowns lead to defects in axons including swellings (black arrows), axon thinning (curved black arrows), MT unbundling (black arrow heads in orange boxes) and gaps in MT bundles (asterisks in blue box); and defects in synaptic terminals including swelling (white arrows) and their breakdown (white arrowheads). Scale bars in A and B represent 10 µm, and 30 µm in coloured boxes.

